# Engineered phages evade the complete defense repertoire of highly phage-resistant MRSA clinical isolates

**DOI:** 10.64898/2026.03.26.714447

**Authors:** Sarah M. Voss, Katharine C. King, Devin J. Hunt, Adam A. Wilson, Bruria Samuel, Orazio R. Bagno, Peter F.W. Sparklin, Bridget Cassata, Joshua W. Modell

## Abstract

Phage therapy is a re-emerging approach for antimicrobial-resistant bacterial infections. However, the narrow host range of most phages remains a major barrier to the success and wider adoption of phage therapy. Although receptor incompatibility is often assumed to define phage-host specificity, we demonstrate that anti-phage defense systems are major determinants of host range in *Staphylococcus aureus*. Using a methicillin-resistant *S. aureus* (MRSA) clinical isolate as a model, we characterized the targeting profiles of its 15 defense systems and, for the first time, generated therapeutic phages that evade the full defense repertoire of a multi-phage-resistant strain. In particular, we show that defense-guided phage recombination is a powerful tool that leverages the modular design of phage genomes to replace targeted with untargeted components. Our holistic approach unveils defense synergies that constrain phage evasion and redundancies that allow the simultaneous evasion of multiple defenses. Finally, we show that an engineered phage cocktail prevents the emergence of phage resistance in the model and a second clinical strain with similar defenses. Our work provides a blueprint for translating our expanding knowledge of defense system identity and mechanism into the rational design of effective, next-generation phage therapeutics.

## Introduction

The growing threat of antimicrobial resistance (AMR) has revitalized interest in bacteriophages (phages) as a therapeutic tool for hard-to-treat bacterial infections^1–3^. Emergency individual applications and a handful of ongoing clinical trials have shown that phage therapy (PT) can be safe, effective, and lifesaving^4–6^. In these cases, phage cocktails are used when available to prevent the emergence of bacterial mutants that are resistant to any single phage^7,8^. However, due to the narrow host range of most phages, many clinical isolates lack even a single effective phage^9–12^. A better understanding of the determinants of host range is therefore necessary to increase the efficacy and promote the wider adoption of PT.

The first, and most studied determinant of host range is the presence of suitable phage receptors on the bacterial surface, which can include surface proteins, cell wall components, flagella, pili, or exopolysaccharides^11^. Following adsorption and injection of genetic material, phages can encounter an assortment of bacterial defense systems (DS) that either directly disrupt the phage lifecycle or trigger an altruistic cell death (abortive infection) that protects uninfected neighbors^13–15^. Computational and functional screens have unveiled hundreds of DS in the pan-immune system of bacteria, with individual genomes harboring an average of 5-6 depending on the species^16–19^. While one study in *Pseudomonas aeruginosa* demonstrated a strain-level correlation between DS abundance and phage resistance^20^, several other studies have suggested that host range is dictated primarily by phage receptors, with DS playing only a marginal role^11,21–23^. In *S. aureus*, most phages bind to peptidoglycan-linked wall teichoic acids (WTA)^24^. Given that WTAs are ubiquitous across *S. aureus* lineages^25^, DS may play a larger role in the host range of this species.

Natural phages have evolved several counter-defense strategies, for example introducing “escaper” mutations in DS targets that decrease recognition or acquiring dedicated DS inhibitors^13,26^. Several studies have shown that phages can also recombine with a cryptic prophage or contaminating phage and replace a targeted component with an untargeted mutant^27–29^. The fitness costs of each evasion strategy are often unclear in mechanistic studies but are relevant when evolving a candidate phage for PT. Furthermore, DS are typically studied in isolation, often in heterologous hosts, leaving a critical gap in our understanding of how native defense repertoires function collectively and how phages might coordinate multiple evasion strategies to overcome them. Prior efforts to engineer phages with enhanced potency or host range have focused exclusively on increasing their affinity for the phage receptor or preventing temperate phages from integrating into the host^30,31^. It therefore remains unknown whether a phage can evolve or be engineered to bypass the ’pan-immune’ repertoire of a multiphage-resistant clinical isolate in order to become useful for PT.

Here we dissect the complete defense landscape of single MRSA clinical isolates to enable rational engineering of therapeutic phages. We show that DS are major determinants of host range across a panel of *S. aureus* clinical isolates. Using a nearly pan-phage-resistant model strain, we establish which phages and genetic loci are targeted by each DS, and we experimentally dissect the efficacy and fitness costs of distinct evasion mechanisms. Equipped with this information, we engineer multiple phages that simultaneously evade all DS within the model. We find that exchanging targeted phage components with untargeted functional analogs via phage recombination is a particularly effective evasion strategy. Our holistic approach reveals interdependent defenses that constrain phage evasion and, conversely, evasion strategies that simultaneously circumvent multiple defenses. When administered as a cocktail, the engineered phages prevent the emergence of phage-resistant mutants and demonstrate efficacy against a second clinical isolate with a similar defense profile.

Together, these findings establish a general blueprint for engineering defense-evading phages for hard-to-treat bacterial infections.

## Results

### *S. aureus* phages are predominantly restricted by intracellular mechanisms

To begin exploring the determinants of phage host range in *S. aureus*, we challenged a panel of 10 clinically isolated *S. aureus* strains with a diverse set of 19 phages, comprising seven siphoviruses, nine myoviruses, and three unsequenced phages (**Fig. 1a, Supplementary Fig. 1a**). Our clinical isolates include seven MRSA strains, two of which are also vancomycin-resistant (VRSA), and spanned five sequence types, including six with the common MRSA sequence types of 5 or 8 (**Table 1**). To probe the degree of lysis for each phage-host combination, we plated serial 10-fold dilutions of phage on lawns of each clinical isolate and calculated the efficiency of plating (EOP) relative to the propagation strain (**Fig. 1b**). Across 190 total combinations, we observed three distinct phenotypes: full infectivity (robust plaques with a < 1 log reduction in EOP), partial infectivity (> 1 log reduction, typically hazy zones of clearance with no single plaques), and no sign of infection (**Supplementary Fig. 1b**). Only 28/190 (15%) combinations demonstrated full infectivity, and only three strains were fully infected by more than one phage, allowing for the development of a diverse therapeutic cocktail. However, for the remaining seven strains, zero or one phage showed full infectivity. We would therefore be unable to produce a phage cocktail for the majority of clinical isolates in our panel.

**Fig. 1:**
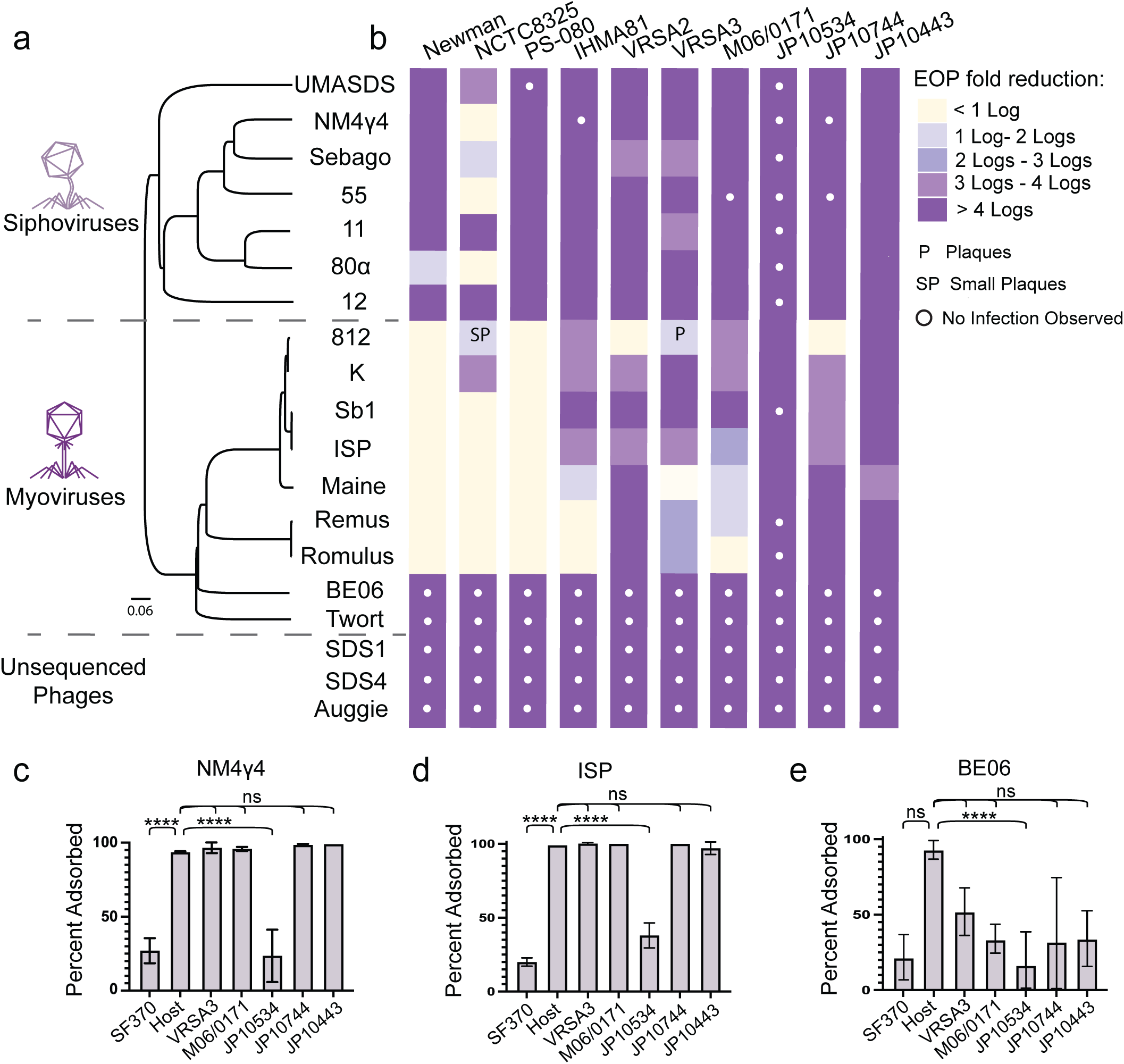
Defense systems, not receptor incompatibility, are the primary determinants of host phage in *S. aureus*. **a,** A phylogenetic tree of phages tested in this study was generated using VipTree based on genome-wide sequence similarities and visualized with FigTree. **b,** The reduction in efficiency of plaquing (EOP) is shown for each phage-host combination relative to the propagation strain for that phage. Beige, full infectivity; white dots, no infectivity, all other squares, partial infectivity. **c-e,** Adsorption assays quantifying the difference in phage PFUs after a 30-minute incubation +/- bacterial cells. NM4γ4 and ISP were broadly able to adsorb, while BE06 was not. Error bars represent SD of n = 2 biological replicates. ****p < 0.0001, ns = not significant.

**Table 1.**

| Strain | Stquence<br>Type | MRSA | VRSA | # of<br>Defense<br>Systems |
| --- | --- | --- | --- | --- |
| Newman | 8 | - | - | 10 |
| NCTC | 8 | - | - | 9 |
| PS-080 | 5 | + | - | 9 |
| IHMA81 | 779 | - | - | 9 |
| VRSA2 | 5 | + | + | 8 |
| VRSA3 | 5 | + | + | 8 |
| M06/0171 | 779 | + | - | 15 |
| JP10534 | 2133 | + | - | 5 |
| JP10744 | 398 | + | - | 9 |
| JP10443 | 8 | + | - | 11 |

We speculated that the 65/190 (34%) combinations exhibiting no infectivity resulted from receptor incompatibility, while the 97/190 (51%) combinations exhibiting partial infectivity indicated intracellular barriers to infection that could be overcome at high phage densities. To test this hypothesis, we performed adsorption assays for one siphovirus and one myovirus that widely demonstrate full or partial infectivity, and an additional myovirus that showed no infectivity on any strain tested. As expected, cases of partial infectivity nearly all showed robust adsorption (>90%), while cases of no infectivity showed greatly reduced adsorption (15-50%) (**Fig. 1c-e**). Notably, one strain and five phages demonstrate broadly reduced adsorption, suggesting that in some cases receptor incompatibility can pose a barrier to PT. However, for the majority of tested strains, intracellular events are the primary determinants of phage host range.

### Abundance of AMR genes and defense systems are correlated

To investigate the extent to which DS contribute to the observed intracellular infection barriers, we used computational tools to identify defense loci in our panel of clinical isolates and in *S. aureus* more broadly. A comprehensive analysis of 13,730 *S. aureus* genomes (including 9,627 MRSA and 16 VRSA isolates) identified 45 distinct DS, with each strain harboring eight on average (**Fig. 2a, Supplementary Fig. 2a**). Of the top 25 DS found in *S. aureus*, 23 were also in the top 25 MRSA DS, and 10 were found among the 11 total DS identified in VRSA (**Fig. 2b, Supplementary Fig. 2b-c**). Over half of these systems (25/45) were found in our panel of clinical isolates, with individual strains carrying between 5 and 15 DS (**Supplementary Fig. 2d**). Notably, the strain with the fewest DS was the same strain that showed broad-spectrum receptor incompatibility (JP10534).

**Fig. 2:**
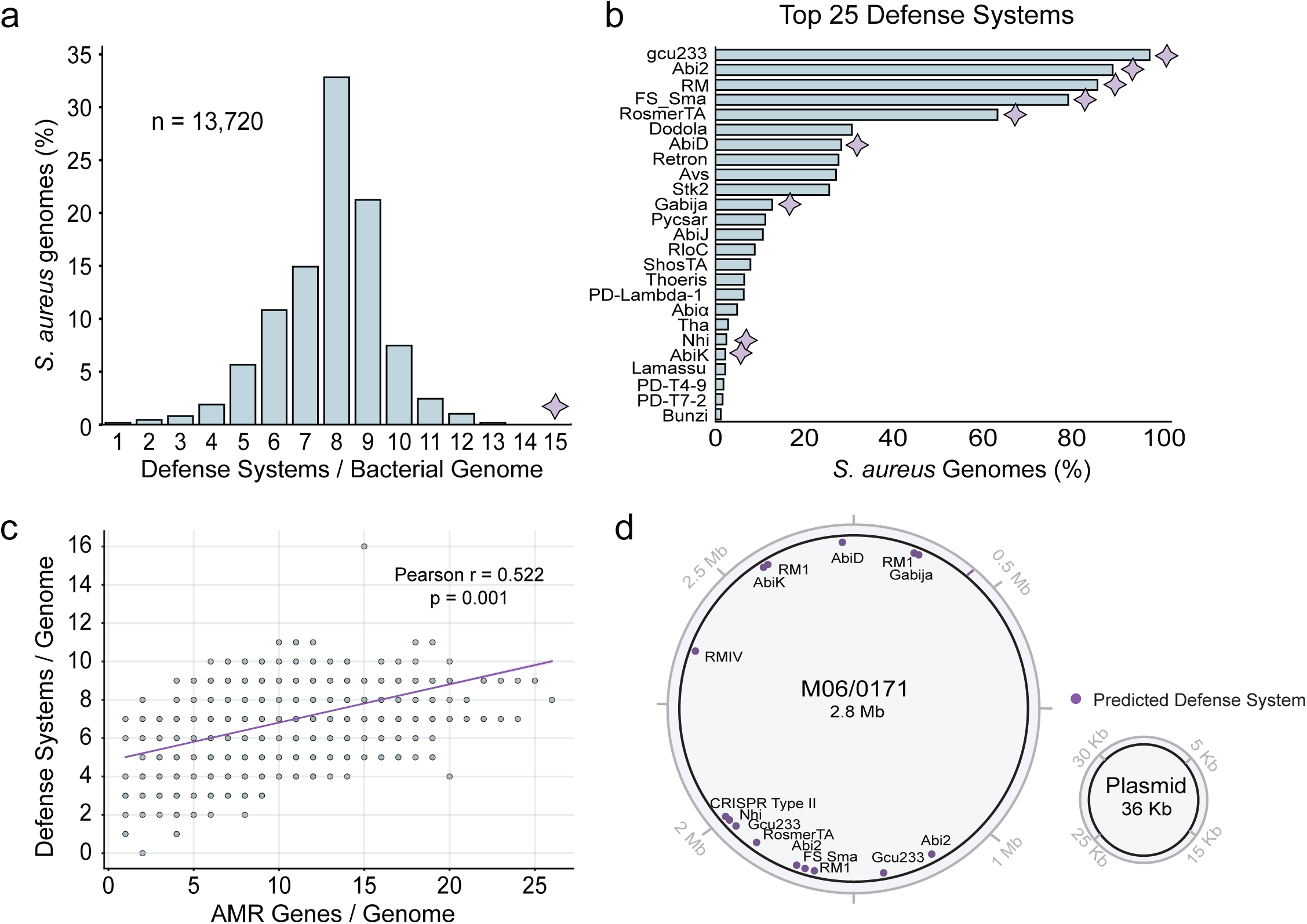
Abundance of defense systems and antimicrobial resistance genes are correlated in *S. aureus*. **a,** Histogram showing the percent of *S. aureus* genomes (n = 13,720 WGS) with the indicated number of predicted DS. The star indicates the number of DS found in the chosen model strain, M06/0171 (M06). **b,** Histogram showing the percentage of *S. aureus* genomes with the indicated DS, for the top 25 most common DS. Stars indicate DS found in M06. **c,** Scatter plot showing the correlation between the presence of AMR and DS genes in *S. aureus* genomes. Fig. S2c shows the number of genomes represented by each dot. **d,** Graphical representation of the genome and plasmid from the high-DS model strain M06, which contains 9 of the top 25 most abundant DS in *S. aureus*. The locations of predicted defense systems (purple dots) and MGEs (purple bars) are shown.

We next examined whether DS abundance correlates with antibiotic resistance, as both DS and AMR genes are known to co-localize on or near mobile genetic elements (MGEs)^32–36^. An analysis of the MRSA genomes revealed a moderate but significant positive correlation (Pearson r = 0.522, p < 0.001) between AMR and DS abundance (**Fig. 2c, Supplementary Fig. 2e**). Overall, this adds to a growing body of evidence suggesting that the antibiotic-resistant strains most in need of phage therapy may also be the most resistant to phage infection^20,34^.

### M06/0171 achieves near pan-phage resistance with a multifaceted defense arsenal

Our empirical and computational results suggest that defense systems may pose a significant barrier to PT in *S. aureus*. For a given clinical strain, we hypothesize that understanding the targeting preferences for the complete defense repertoire will be necessary to develop phages that evade these defenses. Specifically, one must understand for each active defense system (i) which phages are killed, (ii) which phage components or activities are targeted, (iii) what strategies might enable a phage to escape targeting, and subsequently (iv) whether the simultaneous evasion of multiple DS is possible.

To validate this framework, we selected the multidrug-resistant M06/0171 (M06) strain as a model for in-depth characterization. M06, isolated from a pediatric hospital in 2006^37^, encodes 15 predicted DS, including 9 of the top 25 in MRSA (**Fig. 2b, d**). To probe the target range for each predicted DS, we cloned each candidate defense locus with its native promoter onto a low-copy plasmid and introduced them into a defenseless laboratory strain (RN4220). EOPs were calculated as before for each defense plasmid compared to an empty vector. Of the 15 predicted DS, eight were active, and only five - Restriction Modification (RM) 1.1, RM 1.2, AbiK, Gabija, and Nhi - were responsible for the majority (78/81) of defense phenotypes (**Fig. 3a, Supplementary Fig. 3a**). The M06 CRISPR-Cas adaptive immune system encodes 12 DNA memories (spacers), but we found no sequence matches for the phages in our panel and consequently none were targeted. In agreement with the strain-level EOP data (**Fig. 1b**), the only phage that was fully infective on M06/0171 (Romulus) was also the only phage not targeted by any DS (**Fig. 3a**).

**Fig. 3:**
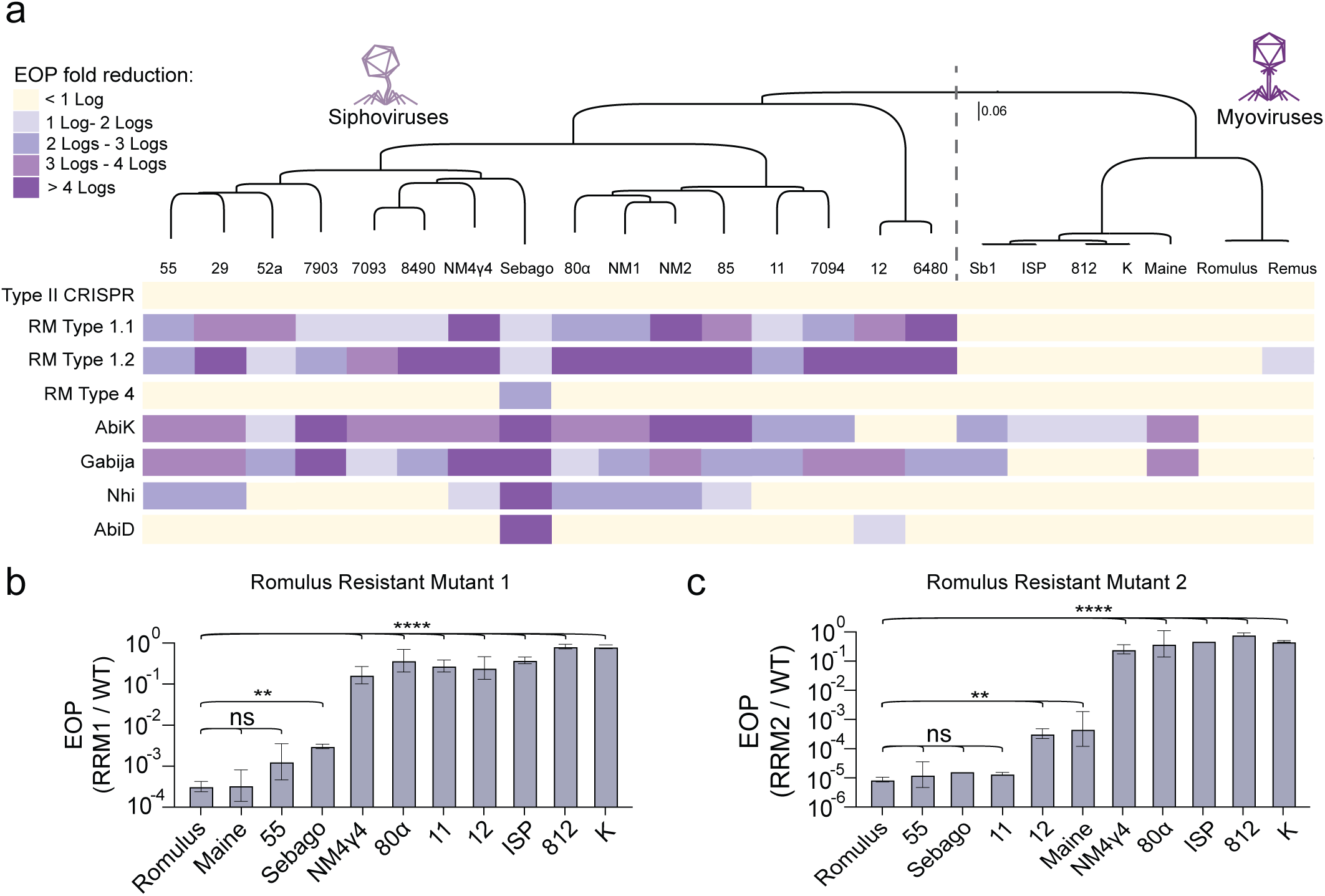
Five active defense systems inform cocktail design against M06/0171. **a,** The reduction in EOP is shown for each phage-DS plasmid combination relative to an empty vector. DS were expressed from their native promoter on a low-copy plasmid in the otherwise defenseless strain RN4220. **b-c,** The EOP is shown for the indicated phage plated on Romulus-resistant mutant 1 (c) or 2 (d) relative to RN4220. Error bars represent SD of n ≥ 2 biological replicates. ***p < 0.001, ****p < 0.0001, ns = not significant.

We sought to complement Romulus in a phage cocktail with phages that can still infect Romulus-resistant mutants. We isolated 2 Romulus-resistant mutants of RN4220 (**Supplementary Fig. 3b, Table 2**) and chose two siphoviruses (80α and NM4γ4) and a myovirus (ISP) that showed full infectivity on both mutants (**Fig. 3b-c**). Next, we sought to understand how to evolve or engineer each phage to overcome the relevant defenses present in M06, beginning with 80α and NM4γ4.

### Siphoviruses can evolve to escape AbiK and Nhi, but not Gabija

The two siphovirus cocktail candidates, 80α and NM4γ4, are each targeted by the same 5 defense systems (RM1.1, RM1.2, AbiK, Gabija, and Nhi) (**Fig. 3a**). Because the likelihood of a phage simultaneously escaping 5 DS is prohibitively low, we sought to isolate escapers for each DS in isolation.

The siphoviruses readily escaped AbiK, as escaper plaques of 80α and NM4γ4 were apparent after infecting AbiK-expressing cells within a top agar lawn. The escaper mutations were located in the single strand annealing protein (SSAP) Sak (80α) or its non-homologous functional analog Sak4 (NM4γ4), which is consistent with AbiK escape mutations found in *Lactococcus lactis*^38^ (**Fig. 4a**). SSAPs are recombination proteins that participate in phage replication by promoting the formation of genome concatemers^39,40^, and while SSAP-mutant escapers could form plaques, they exhibited fitness defects in the absence of defense, limiting their therapeutic potential (**Fig. 4b**).

**Fig. 4:**
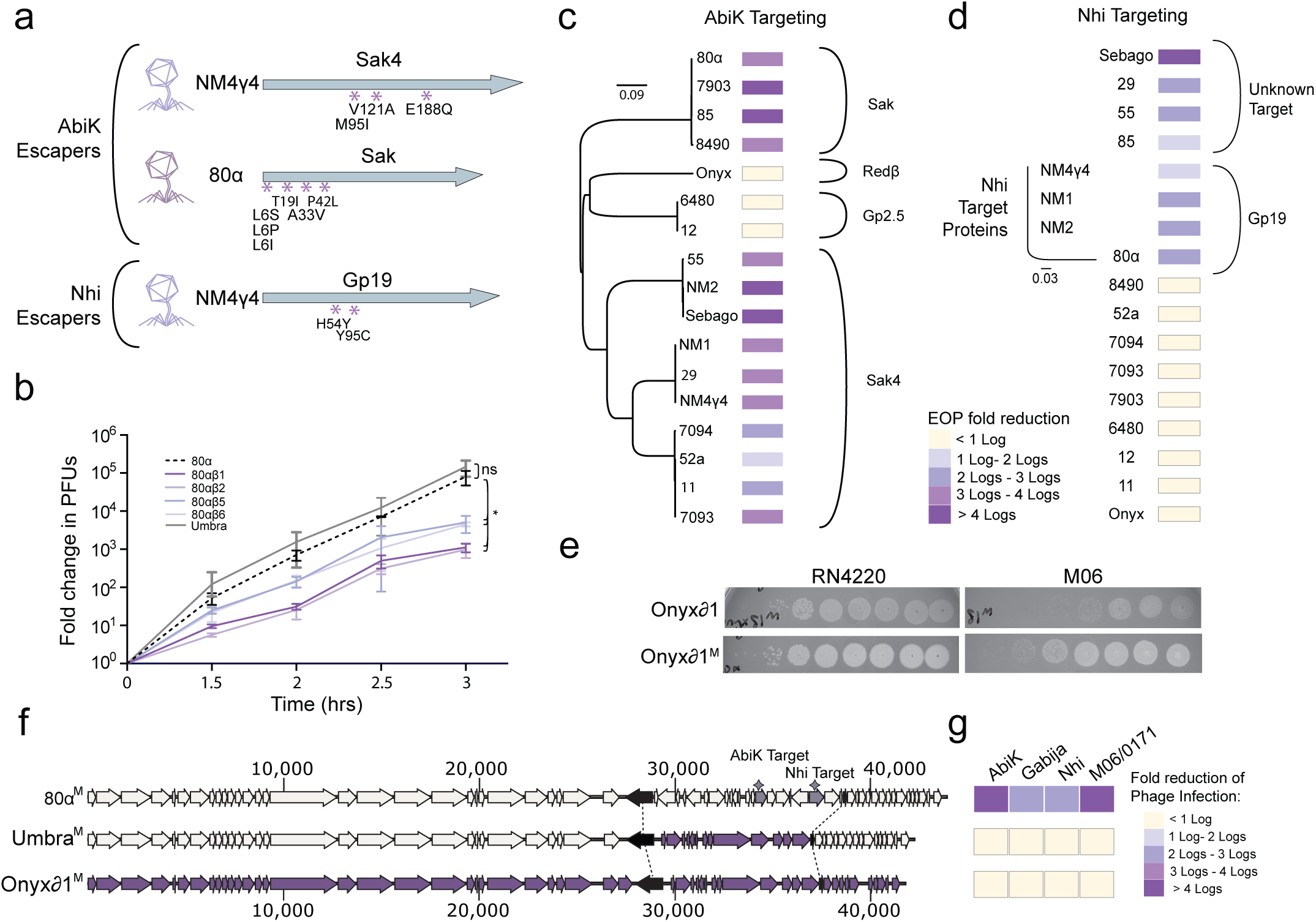
Defense-informed recombination generates novel siphoviruses that evade M06/0171 defense systems. **a,** AbiK escape mutations in 80α and NM4γ4 SSAPs and Nhi escape mutations in NM4γ4 *gp19*. **b,** Time course of parent and escaper phages infecting a defenseless strain, with PFUs at each time point normalized to T=0. Error bars represent SD of n = 3 biological replicates. * p < 0.05, ns = not significant. **c** Phylogenetic tree showing relationships between the SSAP targets of AbiK generated from pairwise sequence similarity with Clustal Omega. The reduction in EOP for each phage on AbiK relative to empty vector is shown in colored boxes. **d,** Phylogenetic tree showing relationships between the targets of Nhi generated as in (c). The reduction in EOP for each phage on Nhi relative to empty vector is shown in colored boxes. **e**, PFU assays of unmethylated and methylated (M) Onyxδ1 on defenseless RN4220 or M06 bacterial lawns. **f,** Depiction of the genomes of 80α, Onyx81, and their recombinant progeny Umbra. Cream represents DNA originating form 80α, purple represents DNA originating form Onyx81, and black indicating genes within which recombination occurred. **G,** The reduction in EOP for each phage in (g) on the indicated DS relative to EV or on M06 relative to RN4220 is shown in colored boxes.

Strikingly, the two phages expressing the SSAP Gp2.5 were not targeted by AbiK at all (**Fig. 3a, 4c**). This reveals a distinct mode of intrinsic immunity: phages carrying non-homologous functional analogs of targeted genes can be inherently invisible to certain defense systems, offering an evasion route that requires no genetic compromise.

Generating Nhi escapers required six passages of NM4γ4 through Nhi-expressing cells. We characterized two escaper plaques, both containing mutations in *gp19*, encoding a replication initiation protein (**Fig. 4a**). A prior study of *Staphylococcus epidermidis* Nhi, which shares 98.7% amino acid identity with M06 Nhi, identified escape mutations in single-stranded binding protein (*ssb*) rather than a *gp19* homolog^29^, suggesting Nhi targets vary across phages. Consistent with this, only 4/8 phages targeted by Nhi (including 80α) contain a homolog of *gp19*, suggesting distinct Nhi targets in the remaining four phages (**Fig. 4d**).

Even after 20 passages of NM4γ4 through cells containing Gabija, no escapers were found. These results underscore the inherent difficulty in modifying the critical activities targeted by Gabija and represent an engineering roadblock.

### An engineered environmental phage evades all active M06 DS

In a recent report, *P. aeruginosa* phages were able to escape Gabija by deleting end-binding proteins, including SSAPs^41^. We wondered whether the universal targeting of the siphoviruses in our panel by Gabija resulted from a lack of SSAP diversity. By searching for new environmental phages in filtered sewage, we isolated four *S. aureus* phages, each having < 95% identity with any previously discovered phage, thus representing unique phage species (**Supplementary Fig. 4a**). Although all four were restricted by M06, one phage (Onyx) formed single escaper plaques at higher phage densities (**Supplementary Fig. 4b**), a phenotype we rarely observed for restricted phages (**Fig. 1b**). Intriguingly, Onyx encodes a member of an SSAP family not previously represented in our phage panel, Redβ (**Fig. 4c**).

We next measured EOPs for Onyx against the individual M06 defenses and found that Onyx was only targeted by the two type I RM systems and not by AbiK, Nhi, or Gabija (**Supplementary Fig. 4c**). We speculated that the escaper plaques on M06 were likely rare phages that acquired protective RM methylation marks before restriction. To generate methylated stocks of Onyx, we passaged it through a strain containing only the methylation subunits of the M06 type I RM systems and found that the methylated phage was no longer targeted by the two active RM systems (**Supplementary Fig. 4d**). Sequencing of the Onyx genome suggested that Onyx is a temperate phage, which are not viable for therapeutic use due to their capacity to mediate horizontal gene transfer^42^. We therefore removed Onyx’s lytic repressor, generating an obligate-lytic variant (Onyx81), and confirmed that Onyxδ1 is no longer capable of lysogeny (**Supplementary Fig. 4e-f**). Finally, we plaqued the methylated, obligate-lytic Onyx81^M^ phage on M06 and observed a reduction in EOP < 1 log, which unlike the parent phage meets the standard for PT^6^ (**Fig. 4e**). Onyx81^M^ was added to Romulus as the second phage in the M06 PT cocktail.

### Phage recombination produces structurally diverse therapeutic phages

Given the possibility that Onyx escapes both AbiK and Gabija with the untargeted SSAP Redβ, we sought to replace the SSAP present in 80α (Sak) with Onyx Redβ using phage recombination. We co-infected cells with Onyxδ1 and 80α and plated on mixed lawns of cells expressing either AbiK or Gabija to select against parental 80α and harboring a spontaneous Onyxδ1-resistant mutation to select against parental Onyx81. We isolated and sequenced the whole genomes of two unique progeny, R4 and Umbra. While R4 was nearly identical to Onyxδ1 (**Supplementary Fig. 4g**-**h**), Umbra was predominantly 80α with only the DNA replication/recombination region from Onyx81, resulting in an exchange of *sak* for *redβ*, and a loss of the Nhi-target *gp19* (**Fig. 4f, Supplementary Fig. 4h**). With these changes, Umbra became resistant to AbiK, Gabija, and Nhi while still expressing all the structural proteins of 80α, resulting in a phage that shared < 95% sequence identity with each parent, thus representing a unique, engineered phage species. Like Onyx, Umbra is partially infective on M06, but when methylated (Umbra^M^) is fully infective (**Supplementary Fig. 4i**). Umbra also inherited the lytic control region from Onyxδ1 and was thus already an obligate lytic phage (**Supplementary Fig. 4j**), joining Romulus and Onyx81^M^ in the cocktail.

Using the same approach as above, we generated six unique recombinant progeny from a co-infection of NM4γ4 and Onyxδ1 (**Supplementary Fig. 4k**). Three of these phages are largely derived from Onyx81, and as expected, escape all the M06 DS and plaque on M06 (**Supplementary Fig. 4l-m**). The other three (Domino, Obsidian, E6), like Umbra, are (i) largely derived from NM4γ4, (ii) have acquired the recombination/replication region from Onyx, and (iii) evade Abik, Gabija, and Nhi (**Supplementary Fig. 4l-m**). However, unlike Umbra, these phages are unable to plaque on M06. We speculate that there is one or more undiscovered M06 DS that restricts these NM4γ4 recombinants.

### Anti-myovirus defense interdependencies show that evasion routes must be considered holistically

We next attempted to engineer a third therapeutic phage, the Myovirus ISP, which was targeted only by AbiK (**Fig. 3a**). After six serial passages of ISP through AbiK-expressing cells, we isolated escapers harboring loss-of-function mutations in *gp075*, a gene of unknown function renamed *gad3* (**Fig. 5a**). A clean deletion of *gad3* (ISPΔgad3) confirmed that the gene product triggers AbiK and is non-essential, and its lack of expression conferring no fitness cost (**Fig. 5b**, **Supplementary Fig. 5a**). Surprisingly, ISPΔgad3 was sensitized to Gabija, which the parent ISP resists (**Fig. 5b**), suggesting that *gad3* encodes a Gabija inhibitor. Plasmid-based expression confirmed that Gad3 inhibits Gabija similarly to the known Gabija inhibitors Gad1 or Gad2 (**Fig. 5c**) despite a lack of homology to either (**Supplementary Fig. 5b**). We attempted to evolve ISPΔgad3 escapers of Gabija but were unable to do so after 20 serial passages, suggesting that ISP cannot simultaneously evade both AbiK and Gabija. This tradeoff illustrates how interdependent defense systems can create evolutionary dead ends that constrain phage engineering (**Fig. 5d**). Future work to uncover the ISP-encoded target of Gabija will be needed to inform successful engineering strategies.

**Fig. 5:**
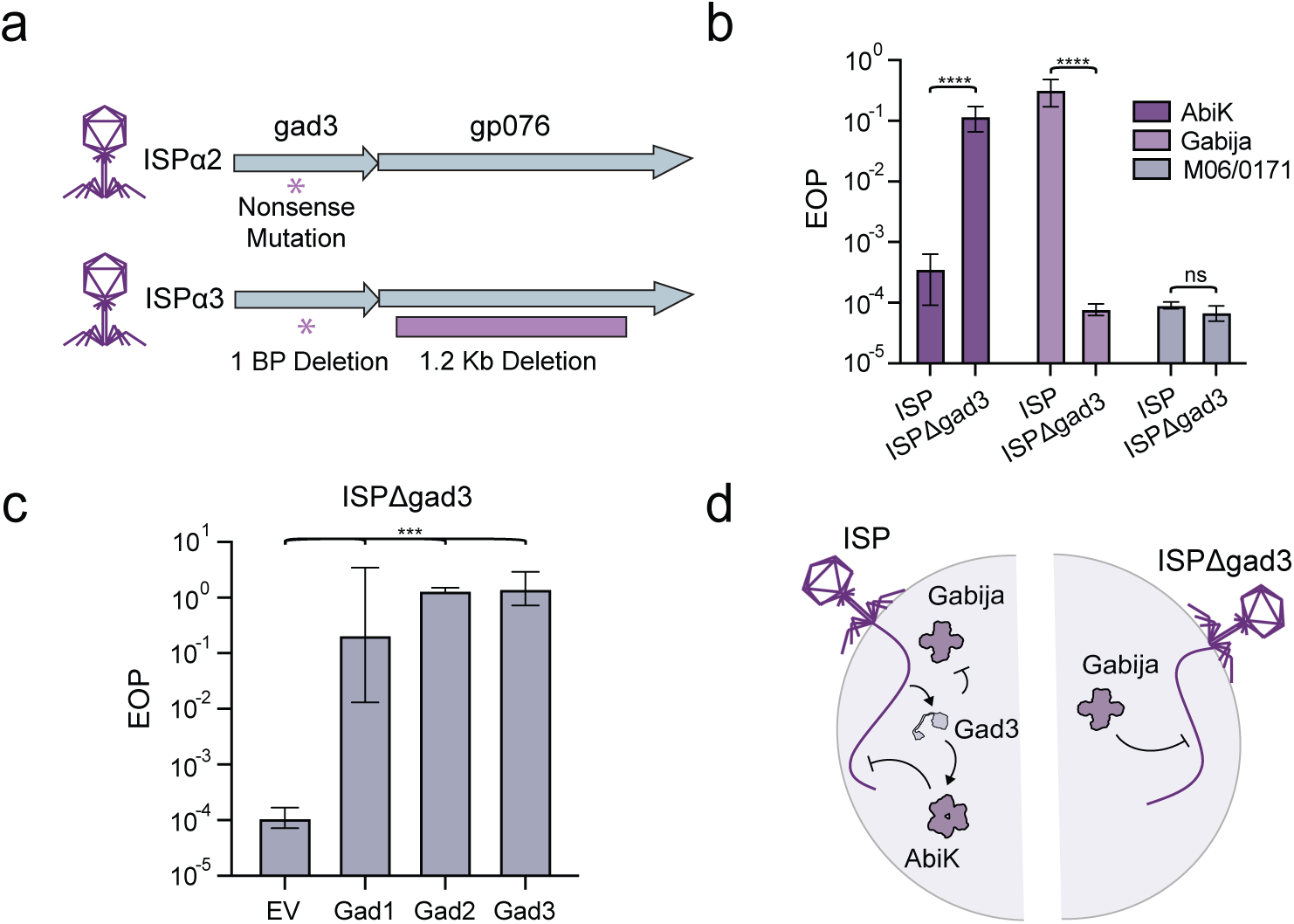
Myoviruses encode a novel anti-Gabija that triggers AbiK. **a,** ISP escapers (ISPα2, α3) emerged after serial passaging on AbiK-expressing cells, each containing loss-of-function mutations in *gad3*. **b,** The reduction in EOP for each indicated phage on cells expressing AbiK or Gabija from a plasmid relative to an empty vector or on M06 relative to RN4220. Error bars represent SD of n ≥ 2 biological replicates. ****p < 0.0001, ns = not significant. **c,** The reduction in EOP of ISPΔgad3 on cells expressing Gabija on one plasmid and the indicated Gabija anti-defense (Gad) gene on a second plasmid relative to cells expressing an empty vector and the indicated second plasmid. Error bars represent SD of n = 3 biological replicates. ***p < 0.001. **d,** Depiction of the evasion trade-off enforced by AbiK and Gabija on phages expressing Gad3. ISP is restricted by AbiK, and while ISPΔgad3 resists AbiK, it becomes sensitive to Gabija.

### An engineered phage cocktail durably kills M06 and a clinical isolate with a similar defense repertoire

With two successfully engineered phages in hand, we asked if Onyxδ1^Μ^ and Umbra^M^ could kill M06 in liquid culture. We infected M06 with engineered or parent phages at an MOI of 1 and monitored cell density for 30 hours. While M06 growth was largely unaffected by the parent phages 80α and Onyx, the engineered phages Onyxδ1^Μ^ and Umbra^M^ lysed the M06 culture within two hours (**Fig. 6a**, **Supplementary Fig. 6a**).

**Fig. 6:**
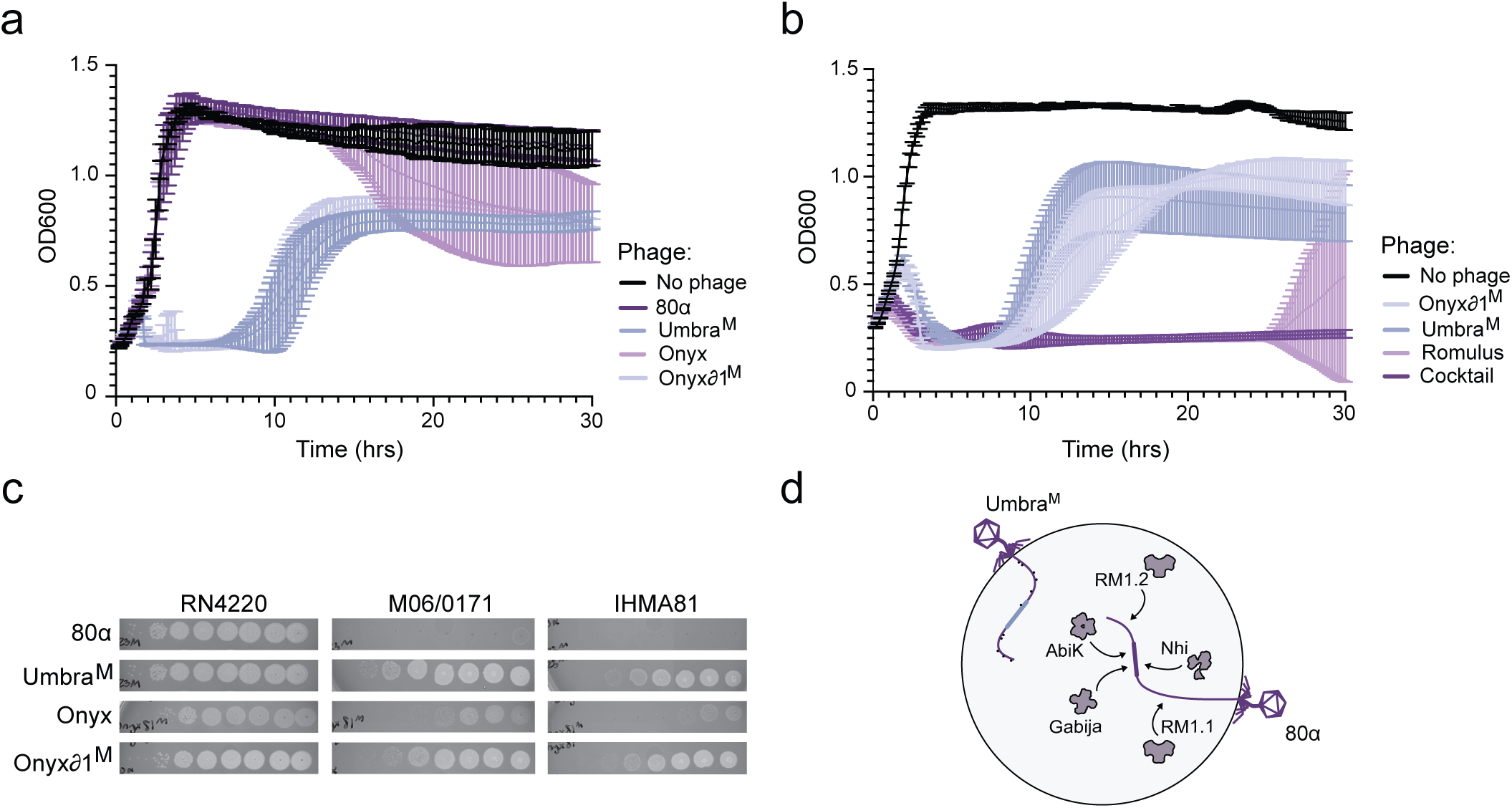
An engineered cocktail durably clears M06/0171 and a clinical isolate with similar defenses. **a-b,** Time course showing the growth in cell density (OD600) of M06 infected with the indicated individual phages or the phage cocktail at MOI = 1. Error bars represent SD of n = 3 biological replicates. **c,** PFU assays showing engineered phages (Umbra^M^, Onyxδ1) and parent phages (80α, Onyx) on the defenseless strain RN4220 or indicated clinical isolates. **d,** Schematic of engineered phage Umbra^M^. The parent phage 80α is restricted by AbiK, Nhi, and Gabija, all of which target the replication/recombination region (thick purple line), as well as two RM systems. Umbra has acquired an untargeted replication/recombination region from Onyx (thick blue line) and methylation marks to evade all five defenses.

However, phage-resistant mutants emerged after ∼10 hours for both engineered phages and at ∼25 hours for Romulus, underscoring the need for diverse phage cocktails.

We next asked whether a three-phage cocktail, consisting of Onyx81^Μ^, Umbra^M^, and Romulus) could prevent the emergence of resistance observed for the individual phages. We infected M06 with each individual phage at MOI 1 or the cocktail at a total MOI of 1. Resistance to individual phages emerged as before (**Fig. 6b**, **Supplementary Fig. 6b**), but M06 cultures treated with the cocktail showed no recovery over 30 hours, confirming phage diversity prevents the emergence of bacterial resistance.

Finally, we tested whether our engineered phages gained the ability to infect other clinical isolates with similar defense systems. IHMA81, like M06, contains AbiK, Nhi, and Gabija, among its 9 predicted defense systems (**Supplementary Fig. 2d**). Given these similarities, we tested Onyx81^Μ^ and Umbra^M^ on IHMA81 lawns and found that they infected several orders of magnitude more efficiently than the parent phages, at or near the therapeutic EOP (**Fig. 6c**). Defense-informed engineering thus offers a new paradigm for therapeutic phage development (**Fig. 6d**), one that yields sustained bacterial suppression with broad activity against clinically relevant strains.

## Discussion

Here, we address a major barrier to the success and wider adoption of PT: for many clinical isolates, few or no effective phages are available. Complex phage cocktails, which prevent the rapid emergence of phage resistance, are even more challenging to assemble. While many studies have pointed to phage receptors as the primary determinants of host range, we show that in many *S. aureus* strains, including MRSA and VRSA isolates, DS are equally or more important. While receptors undoubtedly play a role, as one of the strains in our panel and several of our phages show broad receptor incompatibility (**Fig. 1b-d**), in species with even more diverse receptors like *E. coli* and *P. aeruginosa*, DS abundance is even greater^43^ suggesting that DS are likely a formidable barrier following receptor recognition.

Although many studies have identified and mechanistically characterized DS within the pan-bacterial defense arsenal^15^, ours is the first to ask whether and how phages can circumvent the complete defense repertoire within a single, high DS-containing strain. Our computational analysis demonstrates that *S. aureus* strains encode 8 DS on average and that multidrug-resistant strains are more likely to harbor higher numbers of DS (**Fig. 2a, c**). We used the MRSA clinical isolate M06 as a model, which encodes 15 DS, the most of any strain in our collection. Strikingly, only five of these DS (RM1.1, RM1.2, AbiK, Nhi, and Gabija) were responsible for most of the defense phenotypes (**Fig. 3a**). Eight DS showed no activity against our phage panel.

Some of these may target phages absent in our panel, such as podoviruses and jumbophages, while others may be false positives from the bioinformatic DS prediction programs (like gcu233 and Abi2 which are unusually prevalent).

While the CRISPR-Cas system in M06 did not target any of the phages in our panel, it is genetically intact, and whether the native system can acquire spacers is under investigation. As the only known adaptive immune system in bacteria, CRISPR-Cas systems could endow the defense arsenal with added flexibility and provide an additional avenue for acquired phage resistance. This provides yet another argument for the development of genetically diverse phage cocktails, such that a single spacer would be less likely to target more than one phage.

A therapeutic phage must evade all the active DS within a given strain using three general strategies: **escape** mutations can enable phage components to avoid DS recognition, phages can **inhibit** DS with anti-DS genes, and phages can **recombine** out a targeted module for an untargeted homolog or functional analog. Of the three evasion strategies, evolving escape mutations is the only approach that is agnostic of the defense mechanism and is often used to discover the defense target. However, the feasibility of this strategy depends on the essentiality of the target. AbiK targets the essential phage recombinases Sak and Sak4, and the escape mutations that evade AbiK reduced phage viability (**Fig. 4a-b**). Additionally, generating escapers often requires multiple passages, and for Gabija, escapers could not be identified at all. Even when a single DS is escapable, the probability of escape decreases with each active DS. The time needed to generate escapers, and their unpredictable fitness, makes evolving phages impractical for the production of therapeutic phages for high-DS-containing strains like M06.

Inhibition is another often-used DS evasion strategy. The list of DS blocked by a phage-encoded inhibitor is growing rapidly, and for some DS, multiple structurally and mechanistically distinct inhibitors have evolved^44^. However, inhibitors can create new vulnerabilities, as AbiK cytotoxicity is triggered by the Gabija inhibitor Gad3 in myoviruses like ISP (**Fig. 5**). While Gad3 loss-of-function mutations avoid AbiK, they render phages susceptible to Gabija. The PARIS defense system is similarly triggered by Ocr^45,46^, an RM inhibitor, but how restricted phages evade such compensatory defenses in the same cell has not been studied. Our results illustrate how two interconnected and difficult-to-escape defense systems significantly constrain evasion, highlighting the importance of studying defense and evasion in the context of native hosts with a full defense repertoire.

The final evasion strategy, introducing an untargeted module by recombination, doesn’t risk the fitness defects due to incomplete escape or mutation of an essential target. AbiK, for example, targets Sak and Sak4 SSAPS, while sparing the gp2.5 and Redβ families. Although the direct exchange of modules would require a precise understanding of the modular boundaries, phages are highly recombinogenic, and our defense-guided approach succeeded without this prior knowledge (**Fig. 4f-g**). Curiously, all three of the non-RM defenses in M06 target the replication/recombination region, allowing Umbra to escape them all with a single recombination event. Given the large number of defense systems that target this phage region, it’s possible many phages are one recombination event away from therapeutic potential^47^.

The considerable progress in the last decade to uncover and characterize novel anti-phage DS has opened the door to the rational design of DS-evading therapeutic phages. Our work demonstrates the viability of DS-guided phage design, and a more comprehensive catalog of defenses and inhibitors will only further enable guided immune evasion. We also show that an understanding of DS targets is critical for selecting or engineering phages with untargeted functional modules. In this regard, DS like AbiK with the flexibility to target a set of structurally unrelated phage proteins (Sak, Sak4, and Gad3), pose an additional challenge^48^. Several groups are developing high-throughput techniques to synthesize phages *in vitro*, sometimes assisted by AI models^49,50^. We believe the mechanistic insights in our work and future work will not only guide rational phage engineering efforts but also provide biological constraints and training data to inform AI-based phage design.

## Acknowledgements

The authors thank Rob Lavigne, Jason Gill, Jean-Paul Pirnay, Jose Penades, and Aiden Coffey for sending phages and bacterial strains. We thank Aravind Iyer for helpful discussions and Erin Goley and Brendan Cormack for critical comments on the manuscript. This work was funded by NIH NIGMS (R35GM142731), the Rita Allen Foundation (90094894), an NSF Graduate Research Fellowship for S.M.V., and an EMBO fellowships for B.S.

## Author Contributions

Experiments were conceived and designed by S.M.V. and J.W.M. Experiments were principally performed by S.M.V. Onyx recombination and progeny characterization was performed by K.C.K. Onyxδ1 was created by D.J.H. Assistance with escaper evolutions and plasmid cloning was provided by A.A.W. Computational analyses were performed by S.M.V. and B.S. Onyx was isolated by O.R.B. Assistance with phage preparation and EOP assays was provided by P.F.W.S. The manuscript was written by S.M.V. and J.W.M.

## Methods

### Strains

All experiments were performed using S. aureus strains RN4220 or M06/0171, except for PFU assays on diverse *S. aureus* strains, which were performed in the strain listed, including Newman, NCTC8325, PS-080, IHMA81, VRSA2, VRSA3, M06/0171, JP3, JP4, or JP5. See Supplementary Table 1 for a complete list of strains.

### Phages

The majority of phages were propagated in RN4220 and stored at 4°C as liquid, 0.2 μm filtered stocks in BHI or phage storage buffer (100mM NaCl, 8mM MgSO4, 50mM Tris-HCl [pH 7.5]). A few phages used (UMASDS, Twort, SDS1, SDS4, and Auggie) have different propagation strains. See Supplementary Table 1 for a complete list of phages and phage hosts strains used in this study.

### Standard Growth conditions

All strains were grown in Brain Heart Infusion (BHI) broth at 37°C with shaking at 220 rpm unless otherwise noted. Adsorption assays were performed in LB. Antibiotics were added at the following concentrations: chloramphenicol at 10 µg ml^−1^, spectinomycin at 250 µg ml^−1^, and erythromycin at 10 µg ml^−1^. For phage assays, cultures were supplemented with 5 mM CaCl_2_.

### Isolation of Phage DNA

Before DNA isolation, phage cultures were concentrated using 100kDa centrifugal filters (Amicon), and resuspended in phage storage buffer (100mM NaCl, 8mM MgSO4, 50mM Tris-HCl [pH 7.5]) or media. Then, 160 µl of phage was incubated with 2 U ml^−1^ DNase I and 10 mg ml^−1^ RNase A for 30 min at 37°C. The sample is then brought to a final concentration of 5 mM EDTA and incubated at 75°C for 5 minutes to denature the DNase and RNase. DNA is then isolated from the sample using the DNeasy Blood and Tissue Kit (Qiagen) proceeding with the kit protocol beginning at proteinase K digestion.

### Soft Agar Assays

Each soft agar plate was mixed to include 6 ml of BHI with 0.75% agarose, 100 µl of overnight *S. aureus* culture, 5 mM CaCl_2_, and antibiotics at concentrations listed above, before being poured onto fresh 1.5% agar BHI plates with the corresponding antibiotic. Plates were set at RT until they solidified.

If soft agar plates were used for EOP calculations or phage titering, 3 µl of 10-fold dilutions of phage were then pipetted on top and left to dry at RT, before being incubated at 37°C overnight. Effective phage titers on any strain were determined by counting the number of plaques at the most dilute concentration they could be observed. When only non-confluent lysis could be observed, plaque counts were assumed to be 10. The titer was then calculated by dividing the number of plaques by the µl of original phage stock that was plated at that dilution (PFU/ µl). EOPs were then calculated by dividing the PFU/ µl calculated on a test strain to those calculated on the phage propagation strain, containing an empty vector (EV) control when applicable.

### Adsorption Assays

Cultures used for adsorption assays were grown overnight in LB and diluted back to an OD600 of 0.3 in 1 ml. Phage was added to an MOI of 1 in all bacterial cultures as well as a control containing no bacteria. After incubation for 30 minutes at 37°C, cultures were centrifuged at 16,000 rpm for 1 min. 100 µl was then taken off the top of each culture and immediately used for 10-fold dilutions which were immediately plated onto freshly made soft agar. Soft agar was incubated overnight and PFUs were calculated as previously described. The percent of phage adsorbed was calculated using the following equation: [(phage titer without cells – phage titer with cells)/ phage titer without cells] * 100.

### Cloning and plasmid construction

See Supplementary Table 1 for the oligos and plasmids used in this study. All plasmids were constructed using Gibson assembly^51^. PCR to generate fragments needed for Gibson assembly was performed using Phusion HF DNA polymerase and 5xPhusion Green Reaction Buffer (Thermo) per manufacturer’s instructions. Gibson assemblies were performed as described previously. Briefly, each PCR product was mixed in a total volume of 5 µl in equimolar amounts and then mixed with 15 µl of Gibson master mix and incubated at 50°C for 1 hour. The reactions were then drop dialyzed on MCE membrane filters, 3.0 µm pores, for 30 min and 5 µl was electroporated into 50 µl of RN4220 competent cells.

### Selection for Romulus Resistant Mutants

A soft agar plate of RN4220 was generated using the above method, with the addition of the phage Romulus to the soft agar mix at MOI 10. Individual colonies were selected and re-struck to single colonies twice to remove residual phage. PFU assays were then used to determine the resistance or susceptibility of these strains to various phages.

### Liquid Phage Evolution

One round of evolution of ISP was done by adding ISP (MOI 0.1) to AbiK expressing cells at an OD600 of 0.01 and incubated for 4 hours, before centrifugation was done to remove the remaining cells (4200 rpm, 10 min, 4°C) and phages were filtered through a 0.2 μm filter. Soft agar assays were done after each evolution to determine the resulting phage titer. If the titer was too low to reach an MOI of 0.1 without exceeding 1/5^th^ of the volume of the culture, parent ISP phage was added to supplement the culture, which was needed for evolutions 2, 3, and 4. After 6 evolutions, single plaques were visible on an AbiK lawn. These single plaques were passaged through top agar twice, before being propagated and sent for whole genome sequencing.

The evolution of NM4γ4 against Nhi was done using the same protocol but used an MOI of 1. NM4γ4 also required 6 evolutions before single plaques were observed.

### Sequencing escaper mutations

The DNA of ISP escapers was isolated as described above and was concentrated using the Genomic Clean and Concentrator (Zymo). Concentrated DNA was sequenced through Plasmidsaurus’s long-read sequencing services.

### Anti-Gabija Testing

Known or predicted Anti-Gabija genes were cloned behind a constitutively active promoter and transformed into EV cells or Gabija-expressing cells, generating a two-plasmid system. The effective titer of a Gabija sensitive phage was determined for each combination of plasmids (Gabija + EV, EV + EV, Gabija + each candidate anti-Gabija, and EV + each candidate anti-Gabija). EOPs were calculated to determine the amount of Gabija mediated phage restriction relative to strains expressing the tested anti-Gabija without Gabija present.

### Targeted Phage Engineering

To add or remove genes from ISP, and to generate Onyxδ1, a plasmid containing the desired edit with 200 bp - 500 bp flanking homology regions was cloned and transformed into RN4220. ISP was propagated on this strain to allow for recombination, and recombinant phages were selected by plating to single plaques on RN4220 containing a Type II CRISPR system of *Streptococcus pyogenes* with a spacer targeting only the WT phage. Thus, single plaques would only form for recombinant phages, or CRISPR escapers. Individual plaques were screened for the correct edit via PCR and linear amplicon sequencing (PlasmidSaurus).

### Soft Agar Phage Variant Selection

80α and NM4γ4 escapers of AbiK did not require evolution, and instead, were isolated from soft agar assays. These assays were performed as above, but with the addition of phage at an MOI of 10 to the soft agar. The next day, single plaques were observed, isolated, and sequenced. Because the target of AbiK in *L. lactis* phages is also present in our *S. aureus* phages (SSAPs), Sanger sequencing was used to sequence only the region of DNA containing these genes.

### Phage Multi-step Growth Assay

RN4220 cultures were diluted to an OD600 of 0.1, grown for 1 hour, and again diluted back to a starting OD600 of 0.1. Each phage (parent and escaper) were individually added to liquid cultures at a starting MOI of 0.001 and grown at 37°C for 3 hours.

Starting at 1.5 hours, 1 ml of culture was removed every 30 min and centrifuged at 6000 rpm for 1 minute. Immediately following centrifugation, phage were serially diluted and plated onto soft agar. The following day, PFUs were enumerated and titers were calculated. Each time point was normalized by the titer of the culture at time 0 hours.

### Isolation of Phage from Sewage

Sewage was collected from the Back River Wastewater Treatment center. Upon entering the lab, sewage was centrifuged (4200 rpm, 10 min, 4°C) and filtered through a 0.2 μm filter. Overnight cultures of selection strains were diluted 1:1000 in 8 ml of BHI and supplemented with 2 ml of filtered sewage. Cells were incubated with sewage for 18 hours, before the culture was centrifuged and filtered, as above. Tenfold dilutions of the resulting phage mixture were then plated on soft agar lawns of the selection strain. Single plaques were passaged twice and amplified.

### Methylation of Phage

M06/0171 contains three Type 1 restriction modification systems, each of which is made up of three subunits: hsdR, hsdS, and hsdM. The hsdS and hsdM subunits of each system were cloned onto different plasmid backbones and transformed into one strain. Because this methylation strain does not contain the hsdR subunit, it cannot restrict phage and only adds the methylation marks needed for phage to escape the corresponding RM systems. Thus, to generate methylated versions of a phage, they were passaged through liquid cultures of this methylation strain as detailed above.

To confirm phages had acquired the required methylation marks, phages were plated on soft agar containing cells expressing each full RM system (including the hsdR subunit) individually. PFUs and EOPs were calculated as described above.

### Defense-guided Phage Engineering

First, we generated selection conditions that would restrict both parent phages, while allowing mixed progeny to propagate. Defense system-targeted phages were restricted using plasmid-expressed defense systems. To restrict Onyx, we generated mutants that are resistant to Onyx infection, as described above. These Onyx resistant mutants were transformed with individual defense systems, each of which was mixed into one soft agar plate to generate our selection conditions.

Phages were co-infected into RN4220 at an MOI of 3, each, and left to amplify at 37°C for ∼ 3 hours. Phages were centrifuged and filtered as previously described, before 10-fold dilutions of the mixture were plated on selection conditions. Single plates were passaged twice to remove background phage, propagated, and sequenced.

### Liquid Growth Curves

Log phase cultures of *S. aureus* in BHI were diluted back to OD600 of 0.1, mixed with phage at MOI of 1, added to a flat-bottom 96-well plate (Greiner 655180), and incubated at 37°C in an Epoch BioTek plate reader with shaking. Measurements of OD600 were recorded every 10 minutes for between 25 to 48 hours.

### Quantification and Statistical Analyses

Basic quantification for cell-based assays can be found in Supplementary Table 2. Statistical analyses and graph generation for cell-based assays were completed in GraphPad Prism v10.6.1 and Python v3.12.12. Statistical significance of adsorption, membrane mutant plaquing, phage growth rate, and Gabija inhibitory activity was determined by ordinary one-way ANOVA with Dunnett’s multiple comparisons test comparing each strain to the phage host, each phage to Romulus, each phage to the parent 80α, and each Gad to EV, respectively. The statistical significance of ISPΔGad3 plaquing compared to the parent ISP was determined by unpaired t-test with Welch’s correction. The values of n, and definitions of significance are listed in figure legends. Formatting of graphs and figures was completed in Adobe Illustrator v23.0.1.

Correlation analysis between AMR gene counts and defense system counts per-genome was performed using both Pearson product-moment correlation (for linear relationships) and Spearman rank correlation (for monotonic relationships) to assess the robustness of associations. Statistical significance was evaluated at α = 0.05. Linear regression analysis was performed to model the relationship between variables.

### Genome Acquisition and Assembly

The assemblies of all *S. aureus* genomes from NCBI whole-genome projects were downloaded on March 14, 2020. After excluding genomes with fewer than 1,000 ORFs, the final dataset comprised 13,730 S. aureus genomes. Bacterial sequence types were determined using the PubMLST^52^ database (pubmlst.org) and the *S. aureus* MLST scheme.

The genome of M06/0171 was sequenced, assembled, and annotated by SeqCenter, using Unicycler^53^ for hybrid assembly of Illumina and Oxford nanopore reads.

Phage genomes that were sequenced with Oxford nanopore technology were assembled with the default parameters of Flye v2.9.2^54^. Phage genomes sequenced with short read sequencing were assembled using Spades v3.13.1^55^. All genomes were annotated using Prokka v1.14.6^56^.

### Defense Systems and AMR Identification

Prokaryotic antiviral defense systems were annotated using command line DefenseFinder V2.0.1^43^ with default parameters. For the M06/0171 genome, an additional defense system (RM IV) was identified by using Padloc v1.1.0^57^. Antimicrobial resistance genes were identified using AMRFinder v3.10.24^58^ to classify strains by resistance profile. MRSA strains were defined by the presence of mecA and/or mecC genes, VRSA strains by the presence of vanA and/or vanB genes, and MSSA strains as those lacking these methicillin and vancomycin resistance determinants. A subset of strains (n = 15) harbored both methicillin and vancomycin resistance genes and were classified as MRSA+VRSA and included in both MRSA and VRSA analyses.

### Phage Taxonomy and Comparison

Sewage phages were run through BLASTn core nucleotide database (core_nt) with search restricted to tailed phages (taxid:2731619). The top three most similar, previously discovered phages were selected for further analysis. These 4 phage genomes were run through VIRIDIC (viridic.icbm.de)^59^ to determine if each phage shared a species (95% threshold) genus (70% threshold). Phage phylogenetic analysis was done using VipTree (genome.jp/viptree) v4.0^60^, which generates a proteomic tree based on genome wise sequence similarities computed by tBLASTx. The resulting newick files were visualized in FigTree (http://tree.bio.ed.ac.uk/software/figtree/) before being imported into illustrator.

### Protein Sequence Analysis

MMseqs2 v13.45111^61^ was used to cluster phage proteins into homologous groups based on sequence similarity. All phage genomes were annotated with Prokka v1.14.6 before protein clustering.

---

**Fig. S1:**
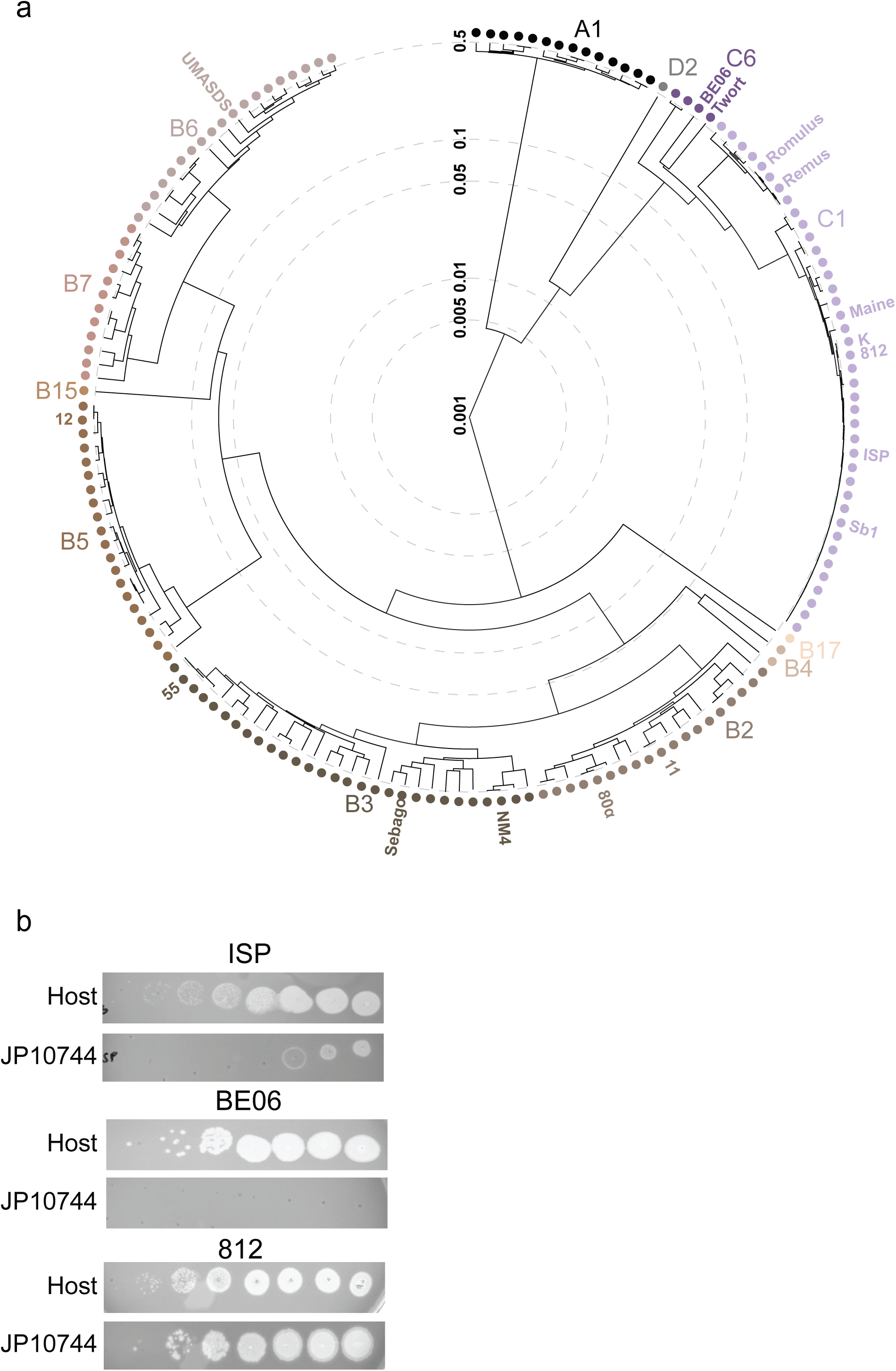
Diversity of phages tested in this study and infectivity phenotypes. **a,** Phylogenetic tree of phages that were clustered by genome-wide sequence similarity in a previous study (REF), supplemented with the phages used in this study. The tree was generated using VipTree which used tBLASTx to compute sequence similarities. **b,** Representative images depicting partial infectivity (top), no infectivity (middle), and full infectivity (bottom), determined by the number of PFUs observed on the propagation strain for that phage (host) compared to the test strain (JP10744).

**Fig S2:**
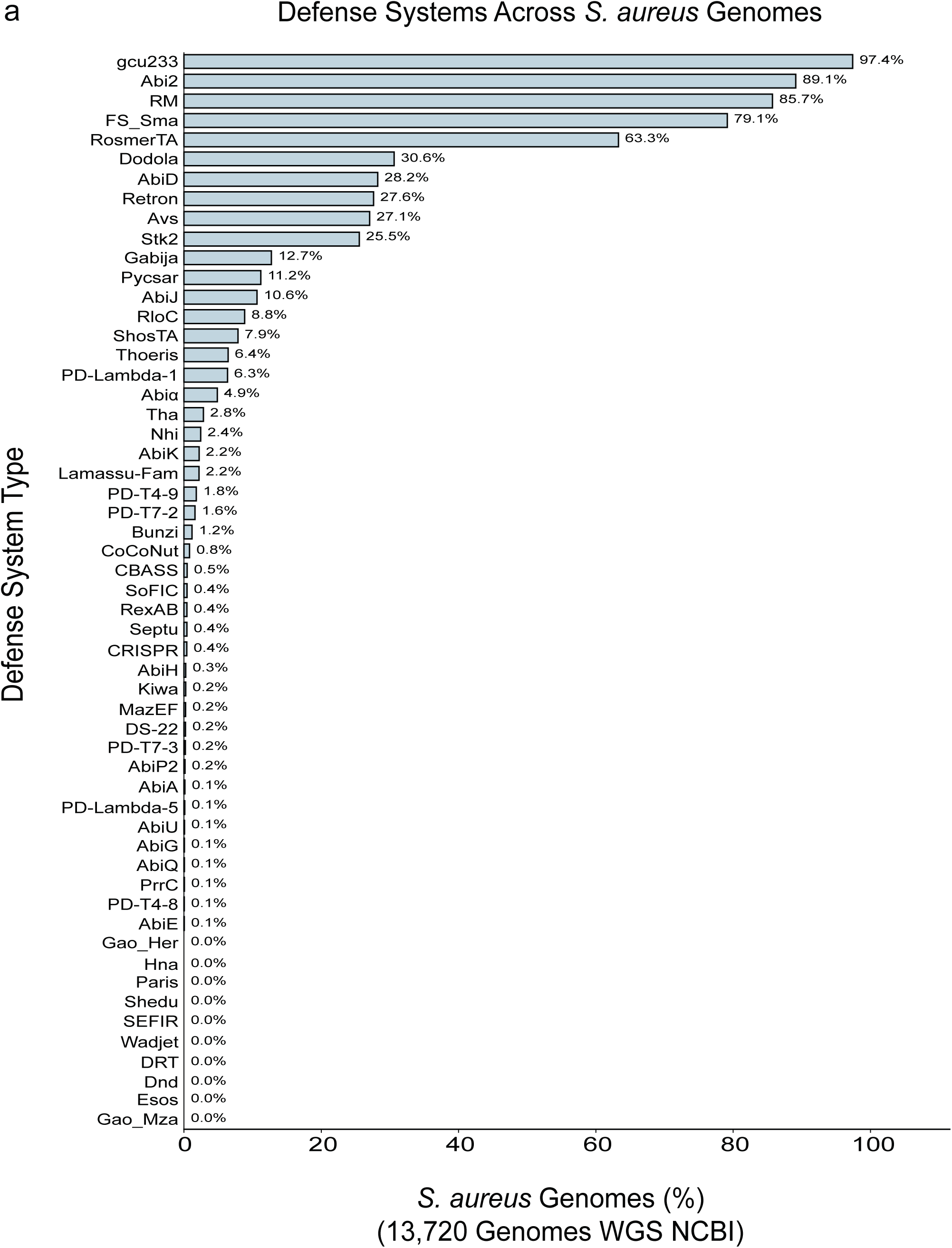
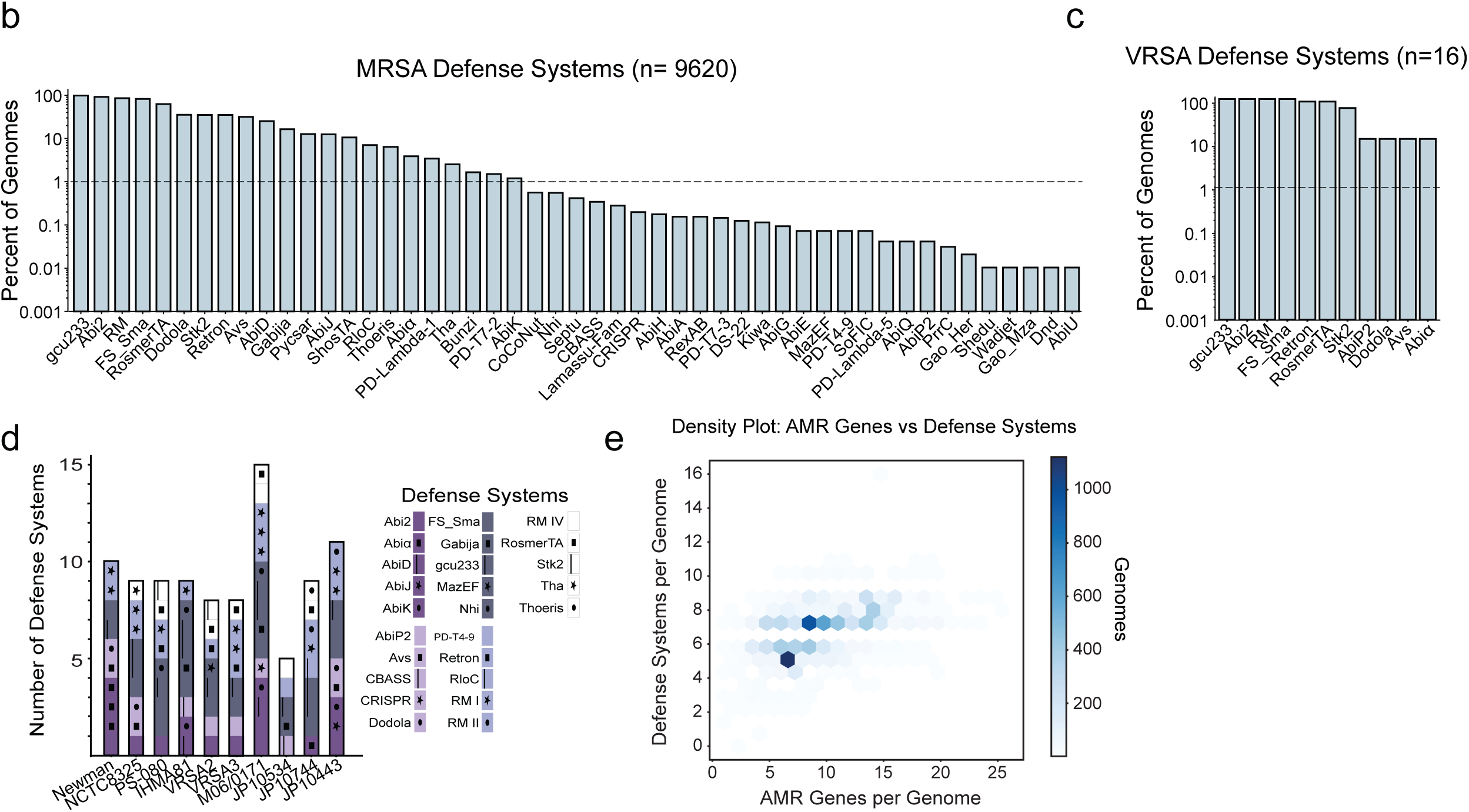
Computational analysis of DS in selected clinical isolates and *S. aureus* more broadly. **a,** DS prevalence across 13,720 *S. aureus* genomes downloaded from NCBI WGS in 2020. DS were identified using DefenseFinder (REFs) and sorted according to prevalence in *S. aureus*. **b-c,** Histogram showing the percent of MRSA (b) and VRSA (c) genomes that contain each DS. **d,** Graph depicting the defense systems present in each clinical isolate from Fig. 1b. Defense systems are listed in alphabetical order and were predicted by DefenseFinder (as well as PADLOC for M06). **e,** Hexagonal binning analysis of 9,627 methicillin-resistant Staphylococcus aureus (MRSA) genomes displaying the density distribution based on antibiotic resistance gene counts and bacterial defense system counts per genome. Color intensity represents the number of genomes within each hexagonal bin, with darker blue regions indicating higher strain density.

**Fig S3:**
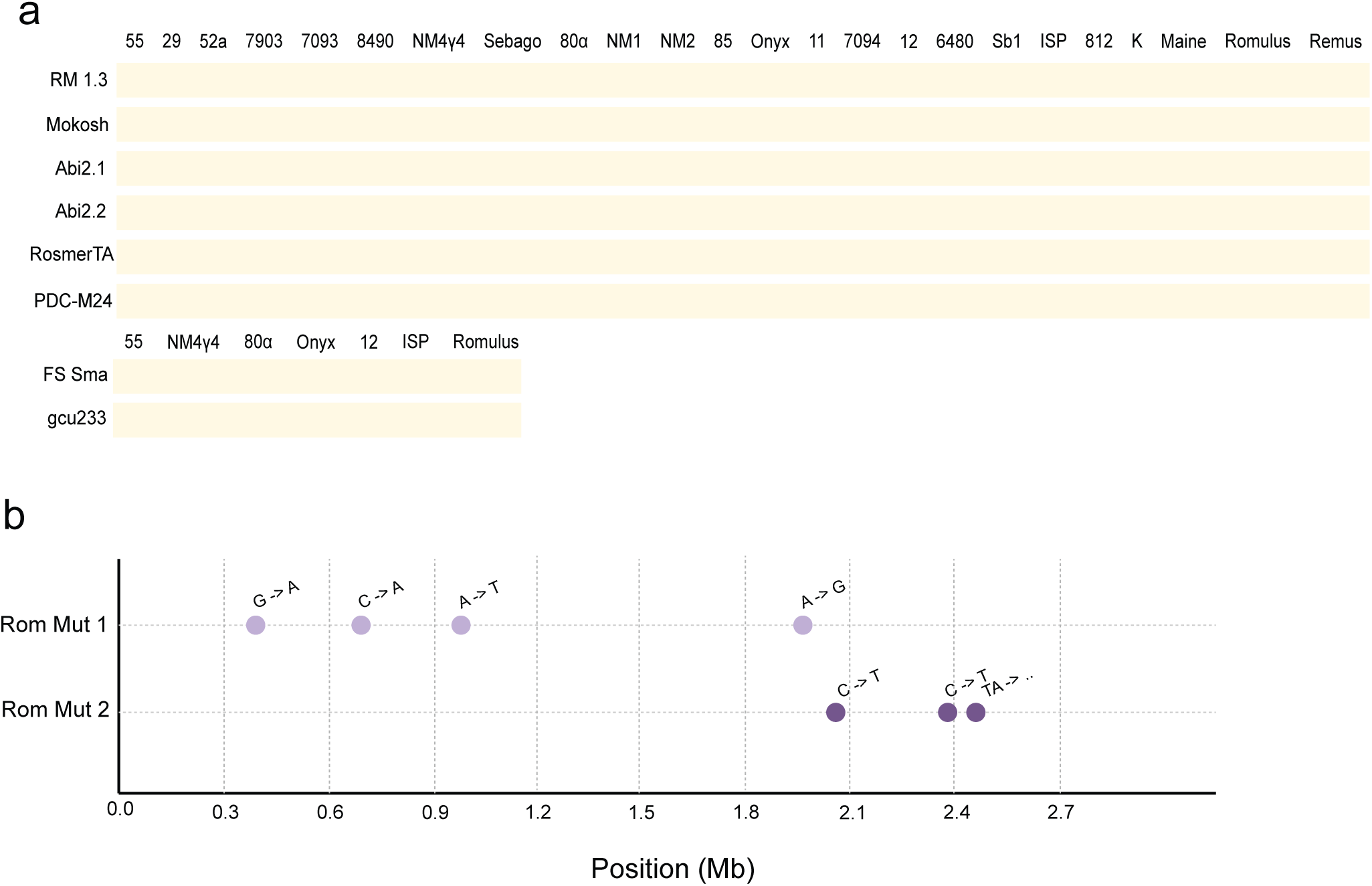
Inactive M06 DS, CRISPR-Cas activity, and Romulus-resistant mutants. **a,** EOPs for each indicated DS relative to EV show no activity against the tested phages. **b,** Graphical representation of the mutations found in two Romulus-resistant mutants derived from RN4220. The diagram was generated using a custom python script.

**Fig S4:**
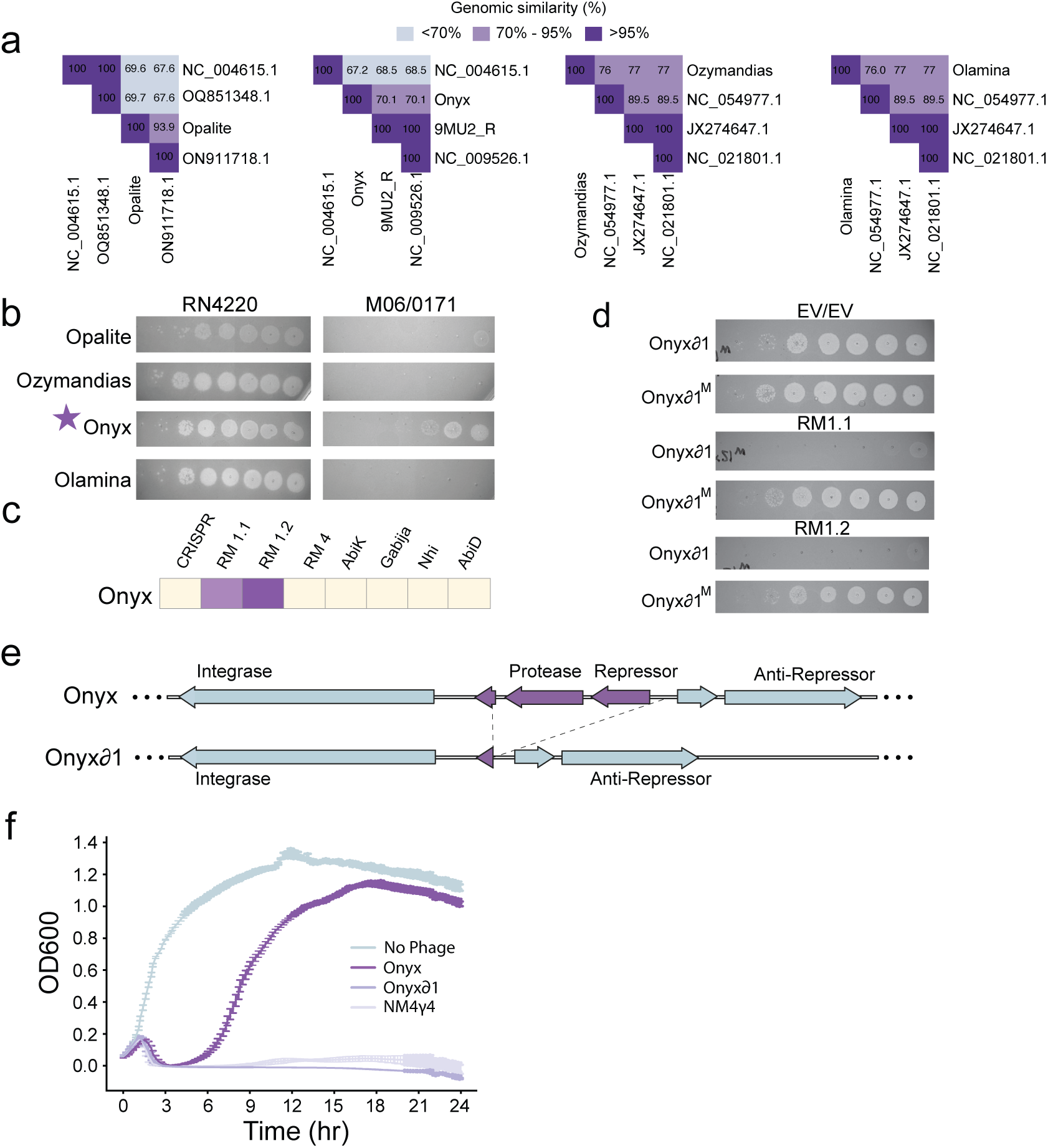
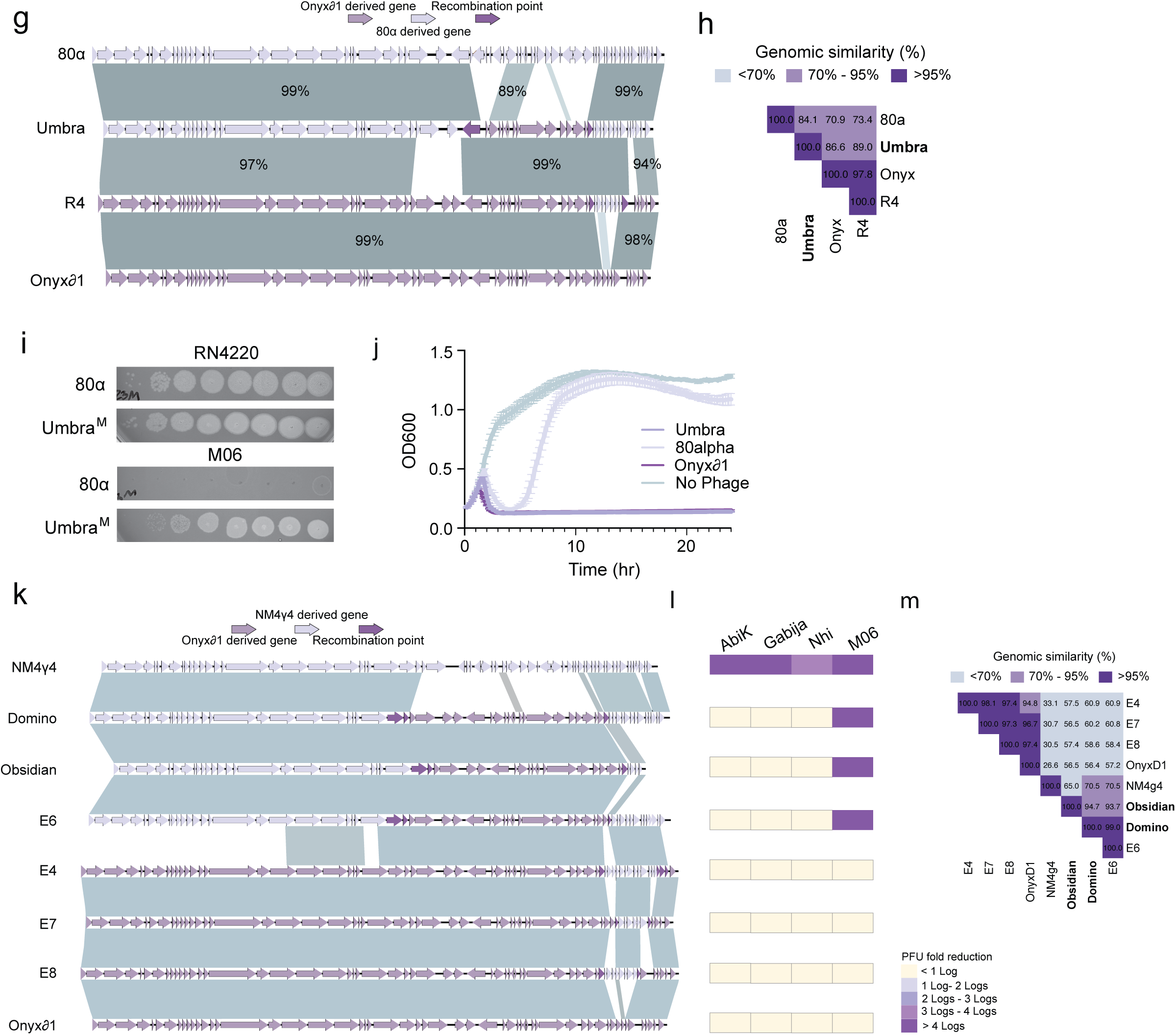
Generation of engineered, M06-killing siphoviruses. **a,** Genomic relatedness of sewage phages to the 4 most similar phages identified in NCBI was assessed using VIRDICT. Recombinants with < 95% average nucleotide identity to any known phages were classified as distinct phage species. All four phages were unique. **b,** PFU assays showing the indicated sewage phages plated on RN4220 or M06. **c,** EOP fold reduction for Onyx on strains expressing the indicated M06 DS relative to EV. **d,** PFUs for unmethylated or methylated (M) Onyxδ1 plated on cells expressing the individual M06 restriction-modification (RM) systems (each subunit expressed on a separate plasmid) or double empty vector. **e,** Graphical depiction of genes deleted to generate the obligate lytic version of Onyx, Onyxδ1. **f,** RN4220 cells were infected with the indicated phages and cell density (OD600) was monitored for 24 hours. **g,** Alignment of parent and progeny genomes, with figures generated using EasyFig (blastn). **h,** Genomic relatedness of recombinant phages to parent phages was assessed using VIRDICT. Recombinants with < 95% average nucleotide identity to both parents were classified as distinct phage species. One novel phage was generated and named Umbra. **i,** PFU assays of the parent phage (80α) and engineered phage (Umbra^M^) on the indicated lawns. Only the engineered phage plaques on M06. **j,** RN4220 cells were infected with the indicated phages and cell density (OD600) was monitored for 24 hours. **k,** Alignment of parent and progeny genomes, with figures generated using EasyFig (blastn). **l,** The fold reduction in EOP on the indicated DS relative to EV or on M06 relative to RN4220 is shown in colored boxes. **m,** Genomic relatedness of recombinant phages to parent phages was assessed using VIRDICT. Recombinants with < 95% average nucleotide identity to both parents were classified as distinct phage species.

**Fig S5:**
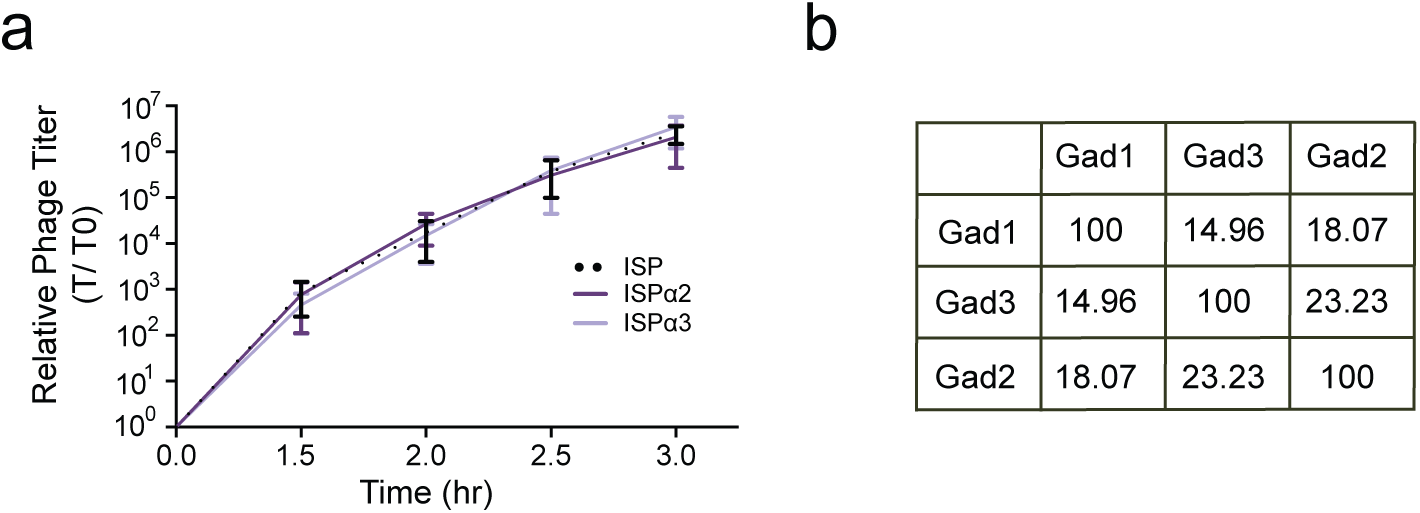
Analysis of anti-Gabija genes found in *S. aureus* myoviruses. **a,** RN4220 cells were infected with ISP or ISP escapers of AbiK at MOI = 0.001, PFUs were measured at the indicated times and fold change was calculated relative to t = 0. All phages show similar replication rates. Error bars represent SD of n = 3 biological replicates. **b,** Percent amino acid identity between the anti-Gabija proteins Gad1 and Gad2, identified previously in *B. subtilis* phages, and Gad3, identified in this study in ISP. Multiple sequence alignment was performed with…

**Fig S6:**
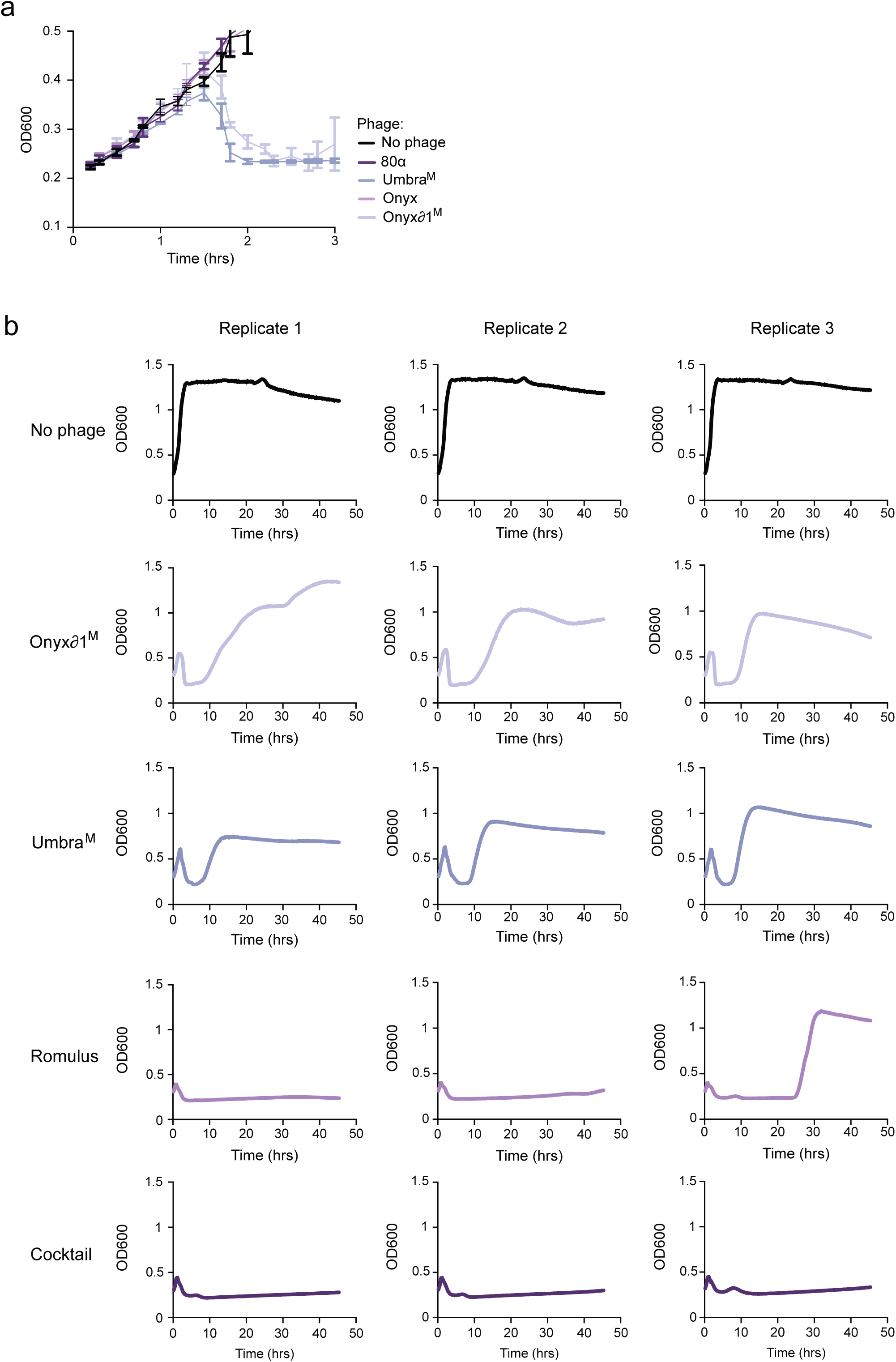
An engineered phage cocktail durably kills M06. **a-b,** M06 cells were infected with the indicated phages at MOI = 1 and cell densities (OD600) were monitored over time. Error bars in (a) represent SD of n = 3 biological replicates. Each graph in (b) shows a single replicate.

